# AA-amyloidosis in cats (*Felis catus*) housed in shelters

**DOI:** 10.1101/2022.05.04.490646

**Authors:** Filippo Ferri, Silvia Ferro, Federico Porporato, Carolina Callegari, Chiara Guglielmetti, Maria Mazza, Marta Ferrero, Chiara Crinò, Enrico Gallo, Michele Drigo, Luigi M. Coppola, Gabriele Gerardi, Tim Paul Schulte, Stefano Ricagno, Monique Vogel, Federico Storni, Martin F. Bachmann, Anne-Cathrine Vogt, Serena Caminito, Giulia Mazzini, Francesca Lavatelli, Giovanni Palladini, Giampaolo Merlini, Eric Zini

**Author notes:** **Corresponding author:** Silvia Ferro, Mailing address: Viale dell’Università, 16 – 35020, Legnaro, Italy.

## Abstract

Systemic AA-amyloidosis is a protein-misfolding disease that is characterized by fibril deposition of serum amyloid-A protein (SAA) in several organs in humans and many animal species. Fibril deposits originate from abnormally high serum levels of SAA during chronic inflammation. In domestic short-hair cats, AA-amyloidosis has only been anecdotally reported and is considered a rare disease. Here we report that an astonishing 57-73% of early deceased short-hair cats kept in three independent shelters suffer from amyloid deposition in the liver, spleen, or kidney. Histopathology and mass spectrometry of post-mortem extracted deposits identified SAA as the major protein source. The duration of stay in the shelters was positively associated with a histological score of AA-amyloidosis (B=0.026, CI95%=0.007-0.046; p=0.010). Presence of SAA fragments in bile secretions raises the possibility of fecal-oral transmission of the disease.

## Introduction

Amyloidoses are classified according to the fibril protein precursor and the distribution of amyloid deposition. More than 30 amyloid proteins have been identified in humans, of which 10 have also been documented in animals [1-3]. AA-amyloidosis, the most commonly reported form in animals, is derived from the deposition of serum amyloid-A protein (SAA), an apolipoprotein produced in the liver upon stimulation by proinflammatory cytokines during inflammatory or neoplastic disorders. This form of AA-amyloidosis is referred to as reactive [4-7]. In companion animals, deposition of AA-amyloid occurs in the familial form in specific breeds, including Abyssinian and Siamese cats and Shar-Pei dogs [1,8].

The exact pathogenesis of AA-amyloidosis has not been fully clarified. However, several hypotheses have been proposed, such as increased circulating concentrations of SAA along with defects in the degrading properties of monocytes or genetic structural abnormalities of proteins [1]. Other factors, such as amyloid-enhancing factors and glycosaminoglycans, may possibly take part in amyloidogenesis by accelerating amyloid-fibril deposition [9].

The transmissibility of AA-amyloidosis has been demonstrated in some animal species as taking place through a prion-like seeding-nucleation-dependent mechanism [10]. In addition, the development of AA-amyloidosis in mice appears to be accelerated by the oral administration of amyloid fibrils extracted from affected humans and bovines, leading to the hypothesis of interspecies transmissibility [11]. The presence of AA-amyloids in feces from cheetahs pointed to a potential oral-fecal transmission route in this species [6].

In animals, the organs most commonly involved are the spleen, liver, and kidneys, as well as gastrointestinal mucosa [5]. Clinical signs depend on the localization and amount of the deposits and reflect the extent of organ damage. Chronic kidney disease is common in affected Abyssinian cats and Shar-Pei dogs, whereas hepatic dysfunction has been reported in Siamese cats with liver deposits. Several studies have described AA-amyloidosis in Siamese and Abyssinian cats, while it has rarely been reported in domestic short-hair cats [12-15]. Amyloidosis was reported in a third of cats naturally infected with feline immunodeficiency virus (FIV), but not in those experimentally infected [16].

The present study was initiated after the observation that 3 consecutive cats from the same shelter were diagnosed with systemic AA-amyloidosis within 6 months. The aims of this study were to investigate the prevalence of AA-amyloidosis in domestic short-hair cats housed in shelters at death, describe clinical and histopathology findings, and assess the possible biliary excretion of SAA fragments.

## Results

### Cats in three independent shelters

Clinical and histopathological data were collected post-mortem from 80 domestic short-hair cats that deceased in three independent shelters 100 km apart from each other in northern Italy. One cat was excluded from the study because sampling was not performed within six hours of death. Twenty-eight (35.4%), 26 (32.9%), and 25 (31.7%) of the investigated domestic short-hair cats were from shelters A, B, and C, respectively. The cats stayed in the shelters for a median duration of 27 months (Q1-Q3: 9-66) and median age at death was 6 years (Q1-Q3: 4-8). Cats were younger in shelter A compared to B (p=0.009) and C (p=0.026), while the duration of stay was shorter in shelter A compared to B (p=0.021), but not to C (p=0.093). Gender distribution was similar among shelters (Table 1). While 12 (15.2%) cats were euthanized due to worsening clinical conditions, 67 cats (84.8%) died spontaneously. Fourteen cats died as a consequence of kidney failure among 47 cats (29.8%) for which the cause of death was reported (Supplementary materials, Table S1). Information on other diseases was obtained only for 22 cats from shelter C; 18 had one or more chronic diseases, the most common of which were feline leukemia virus, chronic enteropathy, gingivostomatitis, and respiratory tract disease (Supplementary materials, Table S2).

**Table 1:** Gender distribution, age and length of stay in the three shelters.

|  | **Females** | **Males** | **Age in years,** | **Length of stay** |
| --- | --- | --- | --- | --- |
| **Cat-shelter** | **N (%)** | **N (%)** | **median (Q1-Q3)** | **in months,**<br>**median (Q1-Q3)** |
| Shelter A | 14 (50) | 14 (50) | 5 (4-6) | 14 (8-28) |
| Shelter B | 10 (38.5) | 16 (61.5) | 7 (5-10) | 38 (17-72) |
| Shelter C | 12 (48) | 13 (52) | 7 (4-10) | 32 (13-84) |

N = number; % = percentage

### Histological and immunofluorescent identification of AA-amyloid deposits in liver, spleen, and kidney

Overall, 38 (48.1%) liver, 46 (58.2%) spleen, and 40 (50.6%) kidney samples of cats (Table 2) had amyloid deposits, with 48 animals (60.8%) showing at least one of the three organs affected (Supplementary materials, Table S3). Of the affected cats, 35 (72.9%) had three concurrently involved organs, 6 (12.5%) had two organs, and 7 (14.6%) had one organ (5 liver, 1 spleen and 1 kidney, respectively). The protein SAA was identified within fibril deposits by immunofluorescence. Custom antibodies raised in mice recognized amyloid deposits in the liver (Figure 1A), kidney (Figure 1B), and spleen (Figure 1C), confirming AA-amyloidosis in all affected cats.

**Table 2:** Histological score of liver, spleen, and kidney samples positive to Congo red staining and showing birefringence under polarized light microscopy in cats.

| **Histological score** | **Kidney** | **Liver** | **Spleen** |
| --- | --- | --- | --- |
|  | **N (%)** | **N (%)** | **N (%)** |
| 0 (negative) | 39 (49.4) | 41 (51.9) | 33 (41.8) |
| 1 (moderate) | 16 (20.2) | 24 (30.4) | 10 (12.7) |
| 2 (marked) | 24 (30.4) | 14 (17.7) | 36 (45.5) |
| Total | 79 (100) | 79 (100) | 79 (100) |

N = number; % = percentage

**Figure 1:**
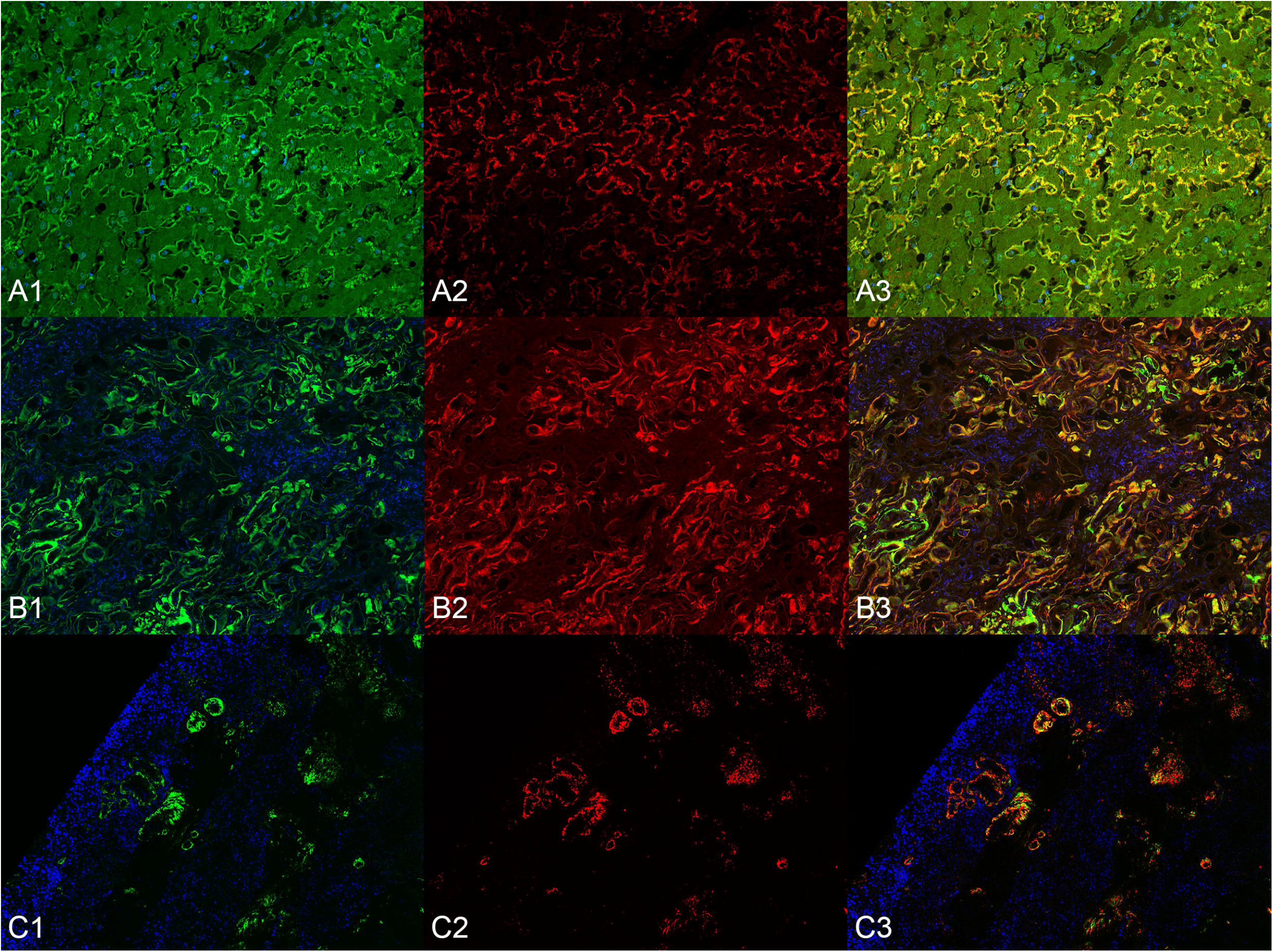
Amyloidosis, cat, fluorescence-immunohistochemistry for AA-amyloidosis. Tissue were stained with Thioflavine S (ThioS) to confirm the presence of amyloidogenic aggregates [A1, B1, C1, left column, ThioS in green, 4’,6-diamidino-2-phenylindole (DAPI) in blue] and with an IgG derived from mice immunized with Qβ-AA peptide vaccine (A2, B2, C2, middle column, IgG in red). A co-localization staining (A3, B3, C3, right column, merge) was generated to authenticate the specificity of the IgGs. A) Liver, magnification 20x. B) Kidney magnification 20x. C) Spleen magnification 10x.

Out of the 48 positive cats for AA amyloidosis, 16 (33.3%), 19 (39.6%), and 13 (27.1%) belonged to shelter A, B, and C, respectively. The post-mortem prevalence of AA-amyloidosis was 57.1% (16/28 cats), 73.0% (19/26), and 52.0% (13/25), respectively. The overall prevalence of AA-amyloidosis as well as distribution of the affected organ was not significantly different between shelters. Of the 22 cats with available information on other diseases, 12 showed AA-amyloidosis in at least one of the three organs with 11 cats (91.7%) being affected by one or more concurrent diseases. The AA-amyloidosis histological score (Table 2) of the spleen was higher than the two other organs (p<0.001). The scores did not differ between liver and kidney, nor between shelters. The distribution of the AA-amyloidosis additive score used in multivariable analysis is reported in Supplementary materials (Supplementary materials, Model S2 and Table S3).

The most common histopathological finding in livers affected by AA-amyloidosis, based on haematoxylin and eosin, was the deposition of abundant eosinophilic material between hepatocytes and sinusoids, causing, when abundant, atrophy and displacement of hepatocytes with derangement of the hepatic cords, which were no longer recognizable (Figure 2). In some cases, there was also a peri-vascular deposition. In the affected spleens, interstitial and peri-vascular eosinophilic deposits were found (Figure 3), often resulting in thickening of the trabeculae. In the affected kidneys, amorphous eosinophilic material in the interstitium and/or glomeruli (Figure 4), multifocal thickening of Bowman’s capsule, atrophy or loss of nephrons, fibrosis, and chronic interstitial nephritis were often noted.

**Figure 2:**
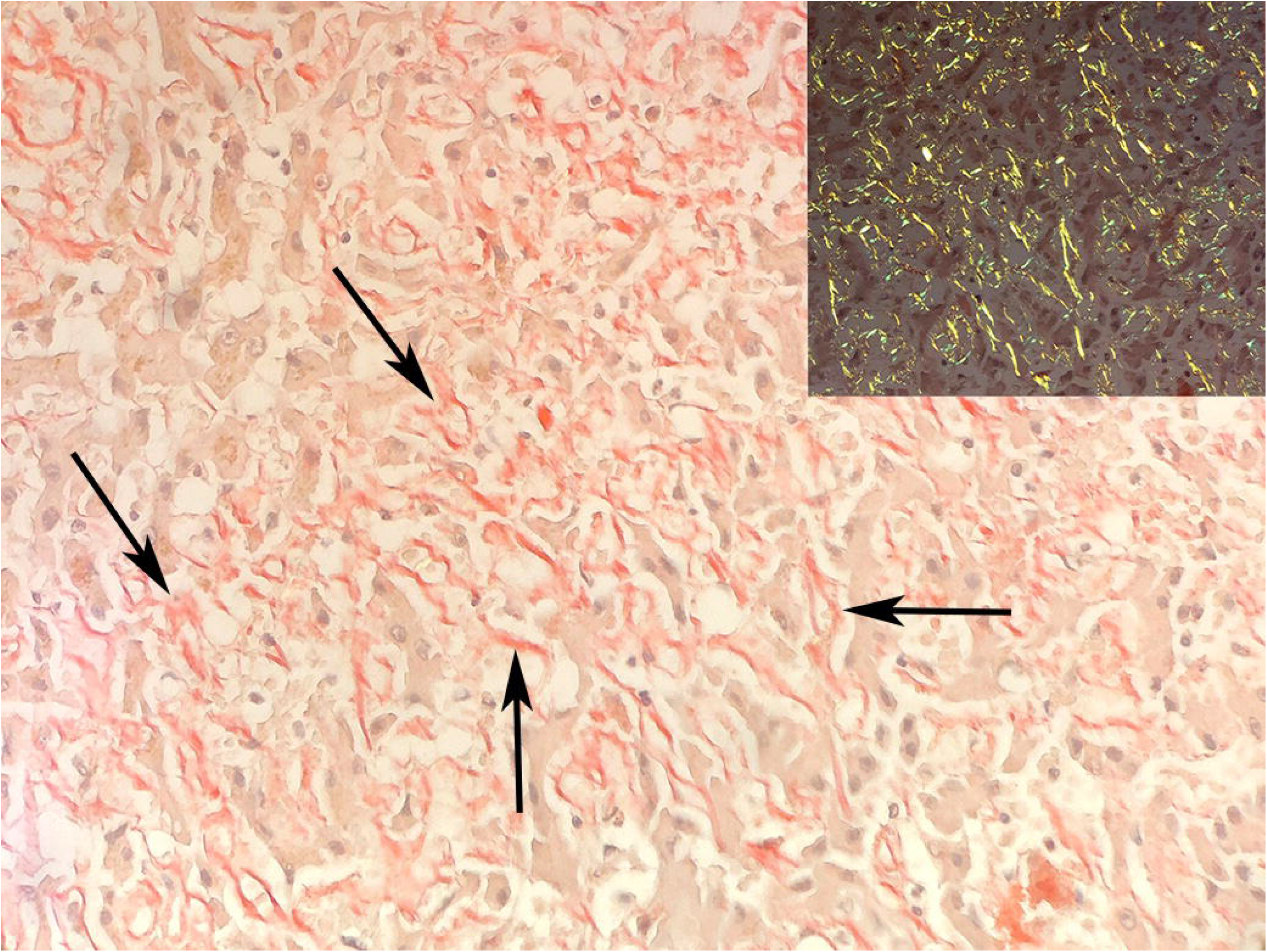
Liver, AA-amylodosis, cat. A diffuse moderate amount of red stained amyloid (arrows) displaces the hepatocellular cords, sometimes isolating the hepatocytes. Magnification 40x. Inset: apple-green birefringence under polarized light. Magnification 20x. Congo red stain.

**Figure 3:**
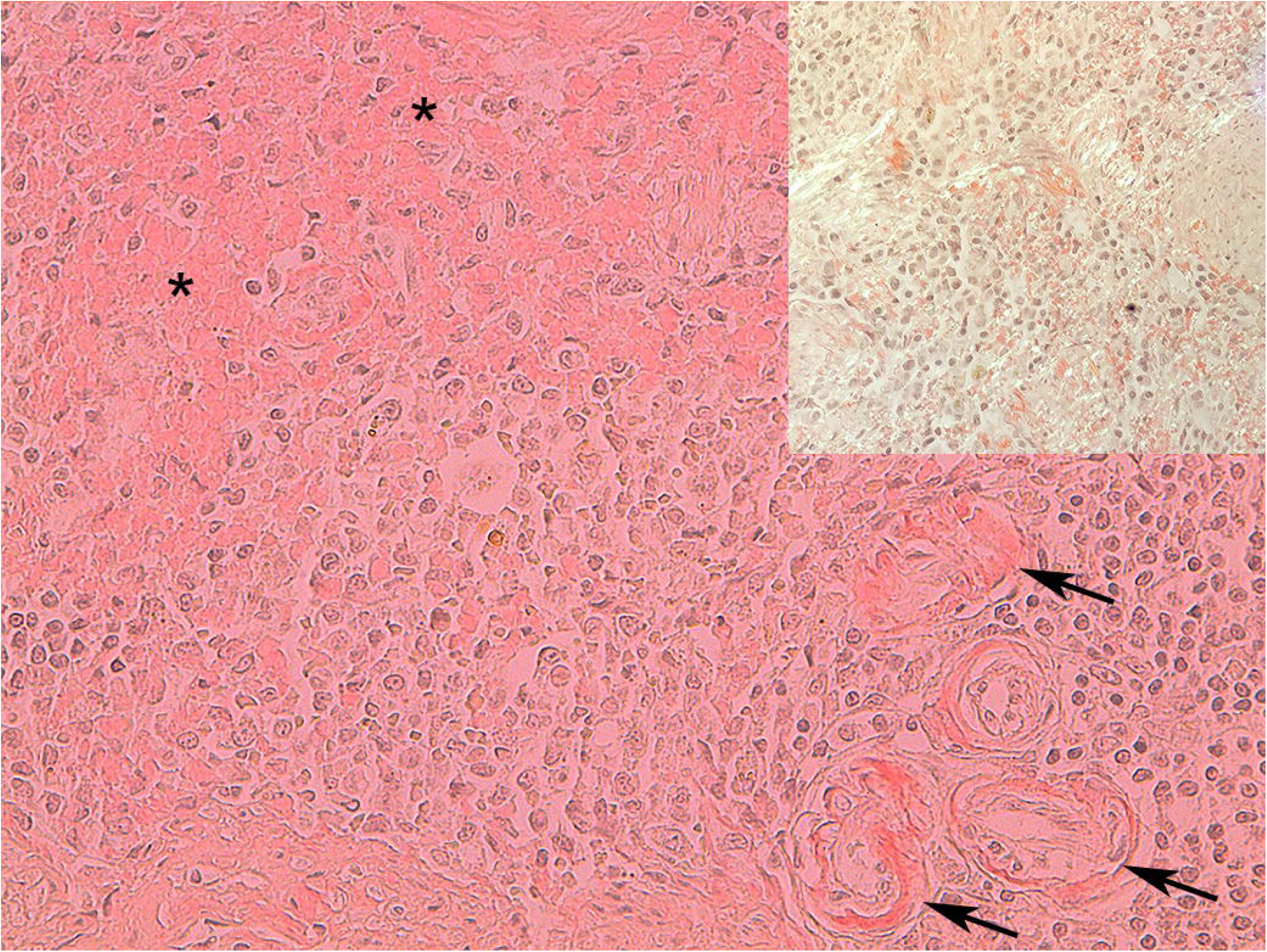
Spleen, AA-amylodosis, cat. Red stained amorphous material (amyloid, asterisks) is observed in the interstitium and around arteries (arrows). Magnification 40x. Inset: apple-green birefringence under polarized light. Magnification 20x. Congo red stain.

**Figure 4:**
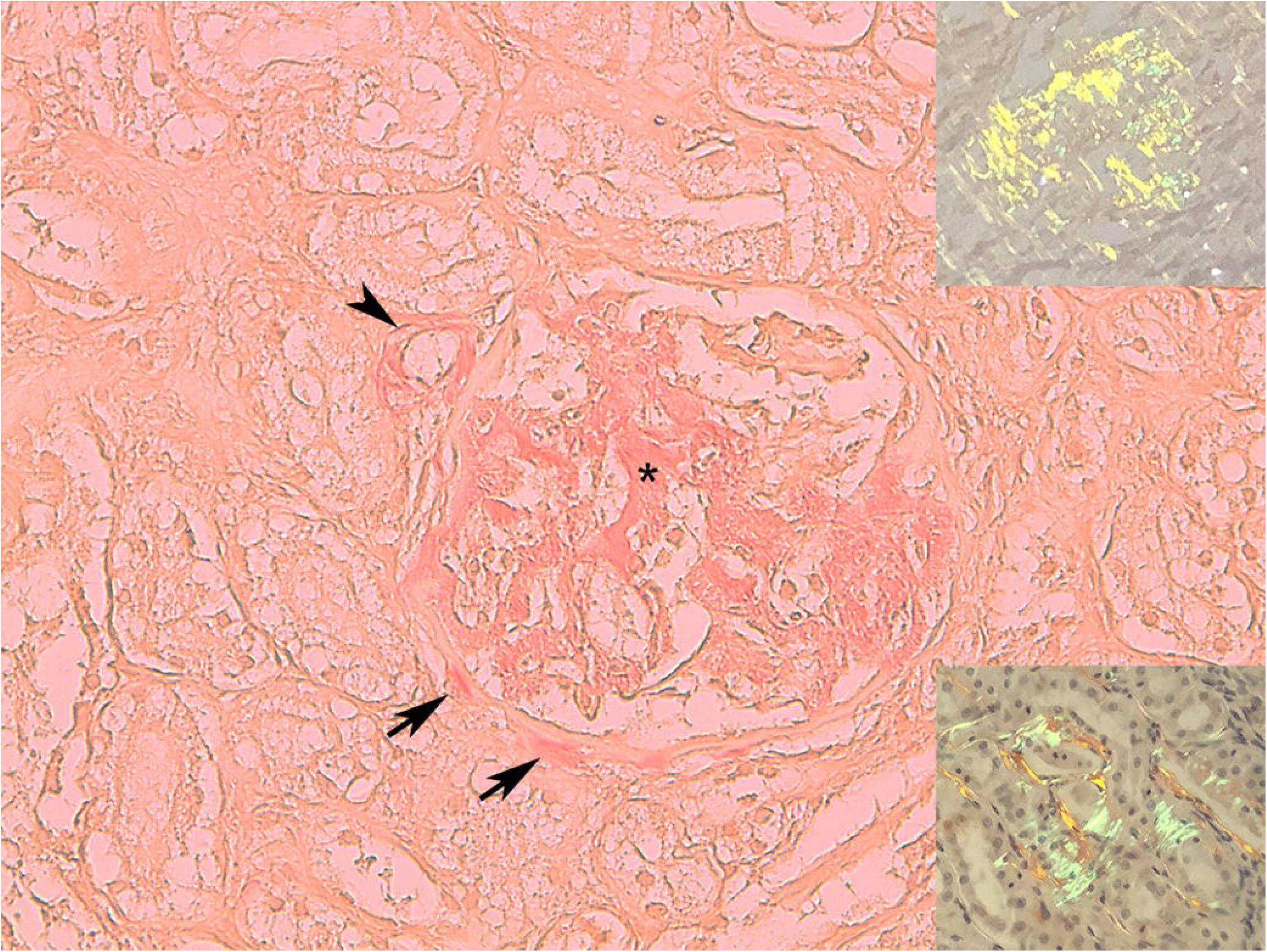
Kidney, AA-amylodosis, cat. The glomerular tuft is expanded by a large amount of Congo red-stained amorphous material (asterisk) consistent with amyloid, which is also present around a proximal arteriole (arrowhead) and along the Bowman’s capsule (arrows). Magnification 40x. Insets: Apple-green birefringence under polarized light is evident in a glomerulus (upper inset) and around tubules (lower inset). Magnification. 20x Congo red stain.

### Presence of SAA fragments in cat bile points to a potential fecal-oral disease transmission route

RT-qPCR analysis of 65 spleen samples from 41 cats with and 24 without AA-amyloidosis were tested for infection with feline immunodeficiency virus (FIV), feline leukemia virus (FeLV), and feline coronavirus (FCoV): three (4.6%), 20 (30.8%) and two (3.1%) cases, respectively, were positive. All FIV (100%), 14 FeLV (70.0%), and one FCoV (50.0%) positive cats had AA-amyloidosis in at least one organ. However, statistical analysis did not reveal any association between any of the three infectious diseases and presence of AA-amyloidosis.

Alternatively, AA amyloidosis may be transmitted horizontally between animals through intestinal shedding of SAA fragments or fibrils. In order to explore this hypothesis, western blot analysis of bile samples from 24 cats revealed that 10 samples from cats with AA-amyloidosis and 5 samples from cats without AA-amyloidosis were SAA-positive, respectively (Figure 5). Two of the 10 amyloid-positive cats with AA-amyloidosis did not have any deposits in the liver. Among the SAA-positive bile samples, the quantity of SAA protein in seven, two, and six samples was rated as mild, moderate, and severe, respectively, based on their intensity levels in western blots. Eight amyloid-positive but none of the amyloid-negative cats excreted bile that was rated moderate or severe (p=0.001). Congo red staining of bile pellets did not detect amyloid fibrils in bile samples, likely due to the low sensitivity of this test.

**Figure 5:**
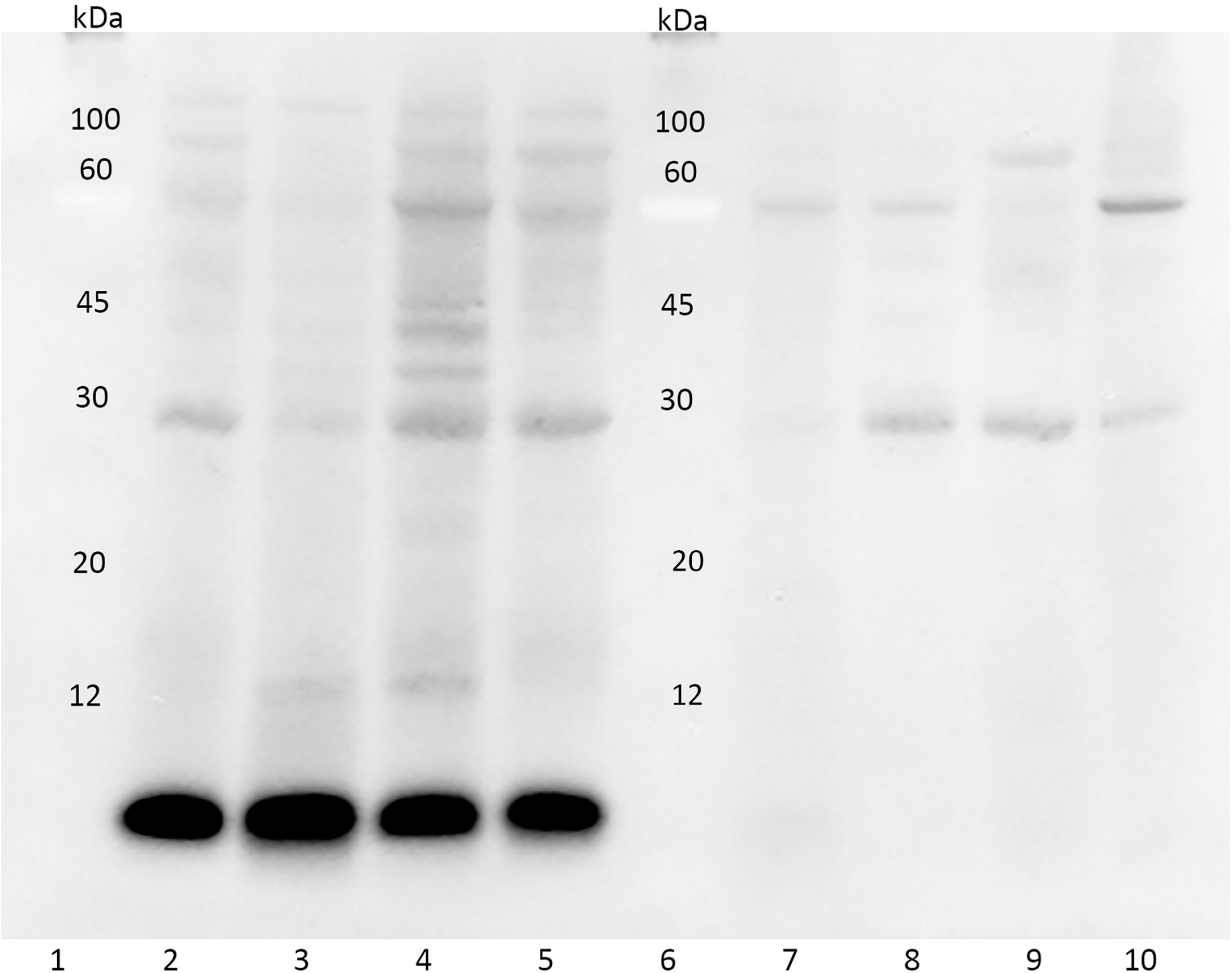
Representative western blot analysis of bile samples with anti-SAA1 polyclonal antibody. Lane 1 and 6: pre-stained molecular weight markers; lane 2-5: positive samples; lane 7-10: negative samples.

### Duration of stay in shelter and duration of other diseases are associated with AA-amyloidosis

Age, gender, and shelter were not associated with AA-amyloidosis nor with the AA-amyloidosis additive score. Duration of stay was not associated with AA-amyloidosis when considered as a dichotomous variable, but was positively associated with the AA-amyloidosis additive score (*B*: 0.026, CI95%: 0.007-0.046; p=0.010) (Supplementary materials, Models S1 and S2).

In the subset of cats in which the disease duration index was calculated, i.e., the proportion of time a cat was sick due to any other disease during its stay in the shelter, those with AA-amyloidosis showed a higher median value compared to those without [0.129 (Q1-Q3: 0.098-0.233) vs. 0.058 (Q1-Q3: 0.006-0.075); p=0.020] (Figure 6).

**Figure 6:**
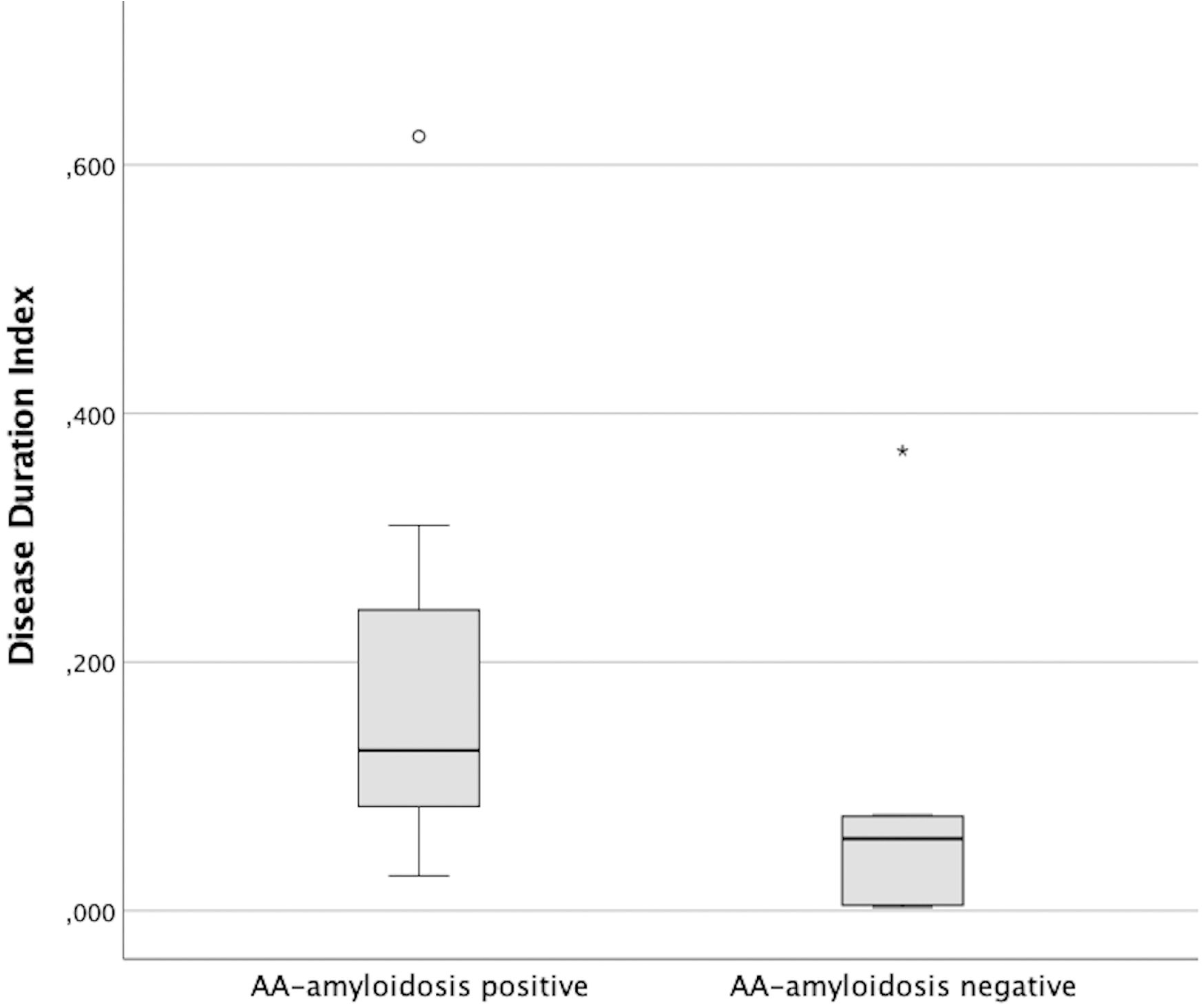
Box and whisker plot of Disease Duration Index in cats with and without AA-amyloidosis.

## Discussion

AA-amyloidosis is the most commonly reported amyloid-related disease in many wild and domestic animals such as mouse, cheetah, and bovine [1]. Although large-scale studies on AA-amyloidosis in cats have not been performed, the disease has been described for Abyssinian and Siamese breeds [1,8]. Few other studies suggested that AA-amyloidosis is a rare disease in domestic short-hair cats [13-15]. Indeed, we previously reported only a single amyloid case among 68 client-owned cats that underwent kidney biopsies, confirming AA-amyloidosis as a marginal disease in short-hair cats [17].

The present investigation puts forward compelling evidence that an almost two-third prevalence of AA-amyloidosis is observed in early deceased cats from three independent shelters. Thus, this study depicts an alarmingly different scenario for cats housed in shelters. Indeed, histopathological post-mortem analysis of 79 cats from three independent shelters, revealed remarkably high AA-amyloidosis prevalence. AA-amyloid deposits were identified in the liver, spleen and kidney by polarized microscopy after Congo red staining and immunofluorescence using SAA-specific antibodies. In a parallel study, we evaluated the structure of amyloid fibrils extracted from the renal tissue of one of the cats from the present study that was deceased from renal failure. This data reveal a cross-β arrangement of wild type cat SAA that is regarded as the defining element of these amyloid deposits [18].

The frequency of AA-amyloidosis was similar in the liver, spleen, and kidney, with more than 70% of cases displaying concurrent deposits in the three organs. Despite the comparable rate, in affected cats deposits were more severe in the spleen compared to the liver and kidney. It is also worth mentioning that in mice the spleen is the organ in which amyloid deposits are most abundant. Macrophages appear to play a key role in the pathogenesis of amyloidosis in the mice spleen [19]. Whether splenic macrophages also contribute to amyloid deposition in the spleen of cats is currently unknown.

The cause of death in most cats with AA-amyloidosis was renal failure associated with amyloid deposits observed in different parts of the kidney. The post-mortem prevalence of AA-amyloidosis was not significantly different among the three independent shelters, and the duration of the stay in shelters was positively associated with the amount of amyloid deposits in affected cats, suggesting a similar constellation of risk factors. Therefore, the exceptionally high prevalence of AA-amyloidosis in shelter-cats may be caused by different factors such as chronic inflammatory diseases, infections, unknown environmental factors, or combinations thereof. In the literature, the deposition of SAA occurred as a consequence of proinflammatory cytokines released during inflammatory or neoplastic disorders [4,5,7]. The vast majority of our cats were affected by concurrent chronic inflammatory conditions, such as chronic enteropathy and gingivostomatitis, which might have triggered amyloid deposition. In addition, the higher disease duration index observed in cats affected by AA-amyloidosis showed that the longer an animal was sick for any reason, the higher the chance of amyloid deposition. Moreover, overcrowding, which is frequent in shelters, might have facilitated the spread of infectious diseases, such as upper respiratory tract diseases, which are common causes of chronic inflammation in shelter cats. The association between AA-amyloidosis with viral diseases, such as natural FIV infection, has been suggested in previous studies [16,20]. However, amyloid deposition was not reported in experimentally FIV-infected cats kept in isolation and without concurrent diseases [16], thus suggesting that the lentiviral infection *per se* is not sufficient for the development of AA-amyloidosis. In this study, FIV, FCoV, and FeLV were not associated with the presence of AA-amyloidosis. In our study, only three cats had FIV, and thus it is possible that by including more cats with FIV an association would have been observed. However, all the above considerations may not be sufficient to explain the extremely different prevalence between client-owned and shelter cats.

Systemic AA-amyloidosis is a well-established case of prion-like disease outside the brain. The transmissibility of AA-amyloidosis has been demonstrated in some animal species through a prion-like mechanism [10, 21-23]. Fecal shedding of AA-amyloid fibrils contributes to disease onset in captive cheetahs [6], and the incidence of AA-amyloidosis is increased by oral ingestion of amyloid fibrils in mice [11]. Orofecal transmission has thus been proposed to play an important role in the development of AA-amyloidosis in captive animals [6,10]. Thus, the possibility of horizontal AA-amyloidosis propagation between cats in shelters should be considered. The presence of SAA fragments in bile proves that they are present in the feces of affected cats. The molecular weight of SAA fragments found in bile samples (Figure 5) is compatible with SAA fragments assembled in SAA fibrils extracted from the renal tissue of an affected shelter cat [18]. Although the presence of SAA fragments in bile samples has been shown herein, its folding, as native or aggregated, could not be established. In any case, bile may be an avenue for SAA fragments to reach the cat intestine and prion-like transmission may take place through feces, as previously shown for other felids [6].

In conclusion, this study demonstrates that the post-mortem prevalence of AA-amyloidosis is remarkably high in domestic short-hair cats housed in shelters, placing AA-amyloidosis among the most relevant health problems for these animals. This observation is in stark contrast with observations in client-owned cats. The time spent in shelters is associated with the amount of AA-amyloid deposits; in particular, the longer an animal was sick for any reason, the higher the chance of amyloid deposition. In this context, we speculate that transmission of the disease may occur through contaminated material ingested by the animal and SAA fragments-containing feces might play a role. Intriguingly, cat shelters may represent a model to investigate the spread of AA-amyloidosis in farm animals as well as in wild animals.

## Methods

### Cats and clinical data

Cats were prospectively enrolled from three shelters in northern Italy, each approximately 100 km apart (shelters A, B, and C). Cats were included if they had died spontaneously or had been euthanized due to the severity of their clinical condition between January and September 2019, and if tissue samples had been collected within six hours from death. Data regarding gender, age at admission to the shelter, length of stay, cause of death, age at death, and presence of concurrent diseases were obtained from medical records.

### Mice and antisera preparation

BALB/c mice (Envigo, Huntingdon, UK) were used for experiments at the age of six weeks and were kept at the central animal facility (Bern, Switzerland). All animals were handled according to protocols approved by the Swiss Federal Veterinary Office (Application #30369). Antigenic preparations from the N-terminal region (MREANYIGAD), the central region (QRGPGGAWAAKV) and the C-terminal region (EWGRSGKDPNHFRP) of feline serum A amyloid protein (SAA) were synthesized and coupled with Qβ virus-like particles (VLPs) [24]. Each VLP-peptide at a concentration of 30 mg/ml was administered once to BALB/c mice. Fourteen days after immunization, mice were bled and the specificity of antisera against each peptide was assessed by ELISA using each synthetic peptide.

### Histology and immunofluorescence

Liver, spleen and kidney samples were collected and fixed in 10% buffered formalin for at least 24 hours and then embedded in paraffin. To identify amyloid deposits, 4 μm thick tissue sections were stained with hematoxylin and eosin (HE) for examination under standard light microscopy, and with Congo red for examination under standard and polarized light microscopy [25]. A semi-quantitative histological score from 0 to 2 was applied based on the amount of Congo red-positive material observed under light microscopy and confirmed under polarized light.

A score of 0 was assigned to samples without positivity, while scores of 1 and 2 were assigned in the case of mild and moderate-to-marked amounts, respectively. To estimate the overall amount of tissue deposits in each affected cat, an amyloidosis score (from 0 to 6) was created by adding the histological score of the liver, spleen and kidney.

Immunofluorescence was performed on cat liver, spleen, and kidney samples to confirm AA-amyloidosis. A total of 1% thioflavine S (T1892, Sigma-Aldrich, Buchs, Switzerland) in ddH2O was used as a stain for amyloid aggregates directly after deparaffinization, and nuclei were stained with DAPI. To examine the interaction of amyloid deposits with antiserum, serum from vaccinated mice (1:10 dilution) was added, followed by a rat anti-mouse IgG conjugated to biotin (Cat#13-4013-85, ThermoFisher, Basel, Switzerland) and a streptavidin conjugated Alexa546 (s11225, Molecular Probes, Eugene, OR, USA). Pictures were acquired with AxioImager A2 and AxioCam (Carl Zeiss, Jena, Germany).

### Infectious diseases in cats

Three slices, 15 μm each, were obtained from formalin-fixed paraffin-embedded splenic samples, and RNA was extracted (High Pure RNA, Roche Diagnostics, Mannheim, Germany). RT-qPCR was used to assess the presence of FIV, feline leukemia virus (FeLV) and feline coronavirus (FCoV). The LightCycler 96 (Roche Diagnostics) was used for amplifications with the Oasig lyophilized OneStep RT-qPCR Master Mix along with three different commercial assays: Feline Immunodeficiency Virus Standard Kit, Feline Leukemia Virus Standard Kit and Feline Coronavirus Advanced Kit (Genesig, Chandler’s Ford, UK). The manufacturers’ instructions were followed for reagents and thermic protocols. Assay performance was assessed by testing 10-fold dilutions of the external positive control (declared concentration of 2×10^5^ viral copies), with a limit of detection (LOD) of 2×10^1^, 2×10^0^, 2×10^0^ and an efficiency of 2.01, 1.87, and 1.88 for the quantification of FIV, FeLV, and FCoV, respectively. Samples were considered positive with Ct<40.

### SAA fragments in cat bile

Samples of bile were collected from the gallbladder of deceased cats. Multiple techniques (transmission electron microscopy, Congo red staining, western blotting) were used to investigate the possible presence of SAA protein or amyloid fibrils in cats’ bile. Briefly, 20 volumes of 0.15 M NaCl were added to 0.15 gm of sample, followed by homogenization for 1 min. After centrifugation at 9418 rcf for 15 min at 20 °C, the supernatants were discarded and pellets dissolved in 20 volumes of 0.15 M NaCl, and then homogenized and centrifuged. This washing step was repeated twice. The pellets were dissolved in 300 μl of ultrapure water, and homogenized and centrifuged at 40,500 rcf for 1 hr at 10 °C. The supernatants were discarded and the pellets were weighed. Ten volumes of ultrapure water were added and, after homogenization, samples were centrifuged at 215,000 rcf for 90 min at 10 °C. Pellets were finally resuspended in 30 μl of ultrapure water. Samples were quantified (Qubit Protein Assay kit, Invitrogen) according to the manufacturer’s instructions. For Western blotting, 30 micrograms of protein extracts were loaded in each well for the analysis. On 24 samples, Laemmli buffer (2x) was added to each protein sample, followed by SDS-PAGE electrophoresis and transferred onto PVDF membranes (Immobilon-P, Millipore, Billerica, MA, USA). Blots were blocked by Tris-buffered saline-bovine serum albumin 1% and incubated with Rabbit pAb SAA1 (AB171030, Abcam, Milan, Italy) diluted 1:1000 in Tris-buffered saline Tween-20 0.1%. Immunodetection was carried out with an alkaline phosphatase-conjugated anti-rabbit immunoglobulin (Vector Laboratories, Peterborough, UK) diluted 1:15000 in Tris-buffered saline Tween-20 0.1% and revealed by a chemiluminescent substrate. The images of the blots were captured by a Chemidoc Touch system (Bio-Rad, Hercules, CA, USA).

Finally, a numerical score was attributed to the results of the western blot, based on the degree of positivity of each sample: 0 for negative samples, 1+, 2+, and 3+ for mild, moderate, and severe positivity, respectively. In addition, the pellets from the bile of 3 cats with liver amyloidosis and displaying positive SAA signal on western blotting were evaluated by Congo red staining, in order to detect possible amyloid fibrils. For Congo red staining, aliquots of the pellet were deposited onto glass slides and dried. Congo red staining was performed using alkaline Congo red–saturated alcoholic solution as previously described [26,27].

### Statistical analysis

Descriptive statistics (median and interquartile interval, i.e., Q1-Q3) were calculated for numerical continuous variables including age, duration of stay in the shelter, and AA-amyloidosis additive score. Frequencies (counts and percentages) were calculated for gender and AA-amyloidosis status (positive or negative), both for the shelter and the entire population of cats. Frequencies of positivity for AA-amyloidosis in the histological samples and for western blot in bile samples were also calculated. Differences between shelters and between cats with and without AA-amyloidosis were analyzed for continuous variables with Kruskall-Wallis and Mann-Whitney tests followed by Bonferroni adjustment. For qualitative variables, rxc contingency tables, chi-square tests with Yate’s correction, or Fisher’s exact tests were used.

In a subset of cats with available medical records and follow-up (n=20, all belonging to shelter C), a disease duration index was calculated arbitrarily. The index represented the proportion of time a cat was sick during its stay in the shelter, irrespective of the disease and severity, and ranged from 0 to 1. The independent sample non-parametric t-test was used to assess the difference in the median disease duration index between cats with and without AA-amyloidosis.

Finally, multivariable analysis that included the relevant predictors for the biological model or the univariate screening with a tolerant p-value <0.15 was used to assess the effects of shelter, age, and duration of stay on the presence of AA-amyloidosis. Two multivariable models were built: a multivariable logistic regression model (Supplementary materials, Model S1) to investigate AA-amyloidosis as a dichotomous variable (positive/negative) and a generalized linear model (Supplementary materials, Model S2) to investigate the severity of AA-amyloidosis deposition based on the AA-amyloidosis additive score. The Hosmer-Lemeshow goodness of fit for Model S1 and Levene’s test of equality of error variance for Model S2, were applied. In both cases, the interaction term between shelter and age or duration of stay was initially included, but excluded from the final models if not significant. The level of significance was set at p=0.05. Statistical analyses were carried out using SPSS v. 26.0 (IBM, Armonk, NY, USA).

## Supporting information

Model S1

Model S2

## Acknowledgments

Valter Fiore (Associazione La Cincia, Val Della Torre, Torino, Italy), Daniela Monfroglio and Davide Pozzi (Associazione Amici dei Gatti Onlus, Galliate, Novara, Italy), Angela Barbato, Federica Folatti, and Francesca Iavazzo are kindly acknowledged for their support.

## Author contributions

F.F., Ch.C., F.P. and E.Z. designed the study and wrote the first draft of the manuscript. F.F. F.P., Ca.C., S.F., S.R., TPS, and E.Z. edited the manuscript. M.D., L.M.C., E.G., and G.G. contributed to the study’s design. S.F., C.G., M.M., M.F., M.V., F.S., M.F.B., A.C.V., S.C., Giu.M. and F.L. performed laboratory investigations. F.F., Ch.C, F.P., Ca.C., G.P., M.D., Gia.M, S.F., and E.Z. analysed data.

## Competing interests

The authors declare no competing interests.

**Correspondence** and requests for materials should be addressed to S.F.

### Institutions where the study was carried out

AniCura Istituto Veterinario Novara, Italy; Istituto Zooprofilattico Sperimentale del Piemonte Liguria e Valle d’Aosta, S.C. Diagnostica Specialistica, Italy; Department of Comparative Biomedicine and Food Sciences, University of Padova, Italy; Department of BioMedical Research, University of Bern, Switzerland; Amyloidosis Research and Treatment Center, Fondazione IRCCS Policlinico San Matteo and University of Pavia, Italy.

### Grant or support

BIRD 193233 University of Padova; 10/19 RC Istituto Zooprofilattico Sperimentale del Piemonte Liguria e Valle d’Aosta; 2019 and 2020 Research Funds AniCura. This study was partially supported by Ricerca Corrente funding from Italian Ministry of Health to IRCCS Policlinico San Donato.

### Meeting where the study was presented

29^th^ ECVIM-CA Congress, Milan (Italy), September 2019.

### Off-label antimicrobial use

none.

### Institutional animal care and use committee (IACUC) or other approval

approved by the Swiss Federal Veterinary Office (Appl. Number 30369)

