## Supplementary material for "AA-amyloidosis in cats (*Felis catus*) housed in shelters": Model S1

**Supplementary materials**

Model S1: Logistic regression model for AA-amyloidosis status (presence vs absence).

|  | OR | Lower bound | Upper bound | P-value |
| --- | --- | --- | --- | --- |
| Age | 0.963 | 0.775 | 1.197 | 0.735 |
| Duration of stay | 1.014 | 0.995 | 1.034 | 0.108 |
| Shelter A | 1.602 | 0.494 | 5.193 | 0.432 |
| Shelter B | 2.604 | 0.784 | 8.654 | 0.118 |
| Shelter C* |  |  |  | 0.295 |

*Shelter C is the reference level. OR, odd ratio.
