## Supplementary material for "AA-amyloidosis in cats (*Felis catus*) housed in shelters": Model S2

**Supplementary materials**

Model S2: Generalized linear model for AA-amyloidosis additive score.

|  | *B* | Lower bound | Upper bound | P-value |
| --- | --- | --- | --- | --- |
| Age | -0.172 | -0.411 | 0.066 | 0.154 |
| Duration of stay | 0.026 | 0.007 | 0.046 | 0.010 |
| Shelter A | 0.225 | -1.079 | 1.530 | 0.732 |
| Shelter B | 0.851 | -0.404 | 2.106 | 0.181 |
| Shelter C* | 0 |  |  |  |

*Shelter C is the reference level. *B*, regression coefficient. R^2^=0.121 (Adjusted R^2^=0.074).
