## Supplementary material for "AA-amyloidosis in cats (*Felis catus*) housed in shelters": Table S1

**Supplementary materials**

Table S1: Cause of death in cats with AA-amyloidosis.

| **Cause of death** | **Number** | **Percentage (%)** |
| --- | --- | --- |
| Kidney failure | 14 | 29.8 |
| Feline infectious peritonitis | 8 | 17.0 |
| Inflammatory bowel disease | 8 | 17.0 |
| Pneumonia | 6 | 12.7 |
| Lymphoma | 3 | 6.5 |
| Seizure due to epilepsy | 3 | 6.5 |
| Uncharacterized anaemia | 2 | 4.2 |
| Hemoabdomen due to hepatic rupture | 2 | 4.2 |
| Bone marrow aplasia | 1 | 2.1 |
