## Supplementary material for "AA-amyloidosis in cats (*Felis catus*) housed in shelters": Table S2

**Supplementary materials**

Table S2: Concurrent diseases in cats with AA-amyloidosis.

| **Concurrent disease** | **Number** |
| --- | --- |
| Feline leukemia virus | 6 |
| Enteropathy | 5 |
| Gingivostomatitis | 5 |
| Respiratory tract disease | 3 |
| Bone marrow aplasia | 2 |
| Cystitis | 2 |
| Toxoplasmosis | 2 |
| Feline infectious peritonitis | 2 |
| Lymphoma | 2 |
| Feline immunodeficiency virus | 2 |
| Ocular carcinoma | 1 |
| Epilepsy | 1 |
| Polycystic kidney disease | 1 |
| Recurrent cutaneous abscesses | 1 |
| Hemotropic mycoplasma | 1 |
