## Supplementary material for "AA-amyloidosis in cats (*Felis catus*) housed in shelters": Table S3

**Supplementary materials**

Table S3: Distribution of the AA-amyloidosis additive score in cats.

| Additive histological score | N (%) |
| --- | --- |
| 0 | 31 (39.2) |
| 1 | 3 (3.8) |
| 2 | 5 (6.3) |
| 3 | 7 (8.9) |
| 4 | 11 (13.9) |
| 5 | 12 (15.2) |
| 6 | 10 (12.7) |
| Total | 79 (100) |

N = number; % = percentage
